# Characterizing neuronal cell bodies in human postmortem cerebral white matter tracts

**DOI:** 10.1101/2025.08.15.670595

**Authors:** Drew VanderBerg, Kelly Perlman, Maria Antonietta Davoli, Gustavo Turecki, Naguib Mechawar

**Affiliations:** McGill Group for Suicide Studies, Douglas Mental Health University Institute, Montreal, Quebec, Canada; Department of Psychiatry, McGill University, Montreal, Canada

**Author notes:** These authors contributed equally (Drew VanderBerg, Kelly Perlman).

**Keywords:** white matter neurons, uncinate fasciculus, corpus callosum, ventromedial prefrontal cortex, human, postmortem, white matter

## Abstract

Until the discovery of white matter neurons (WMN) in the 19th century, white matter (WM) was considered to be completely devoid of neuronal cell bodies. Despite evidence consistently showing evident neuronal soma within cortical WM and their purported implication in neuropsychiatric disorders, these neurons are understudied and have not been characterized in human long-range WM tracts. Using postmortem human brain tissue, we investigated the presence, densities and proportions of excitatory/inhibitory neurons in the uncinate fasciculus (UF) and corpus callosum (CC). We also investigated the ventromedial prefrontal cortex (vmPFC) to validate our methods by comparing our results with previously reported densities of neurons in cortical WM. To identify WMN, we employed fluorescence in situ hybridization with excitatory (SLC17A7) and inhibitory (GAD1) neuronal markers and subsequently validated these neurons at the protein level with NeuN immunohistochemistry. We found that the density of WMN in the vmPFC corresponded with previous independent estimates. The UF displayed a similar, though slightly lower density of WMN compared to the vmPFC, while the CC had a far lower density of WMN than both of these regions. Due to the higher-than-expected density of WMN in the UF, we validated the findings at a second location along the UF temporal segment and confirmed the presence of substantial numbers of WMN in this tract. This research constitutes the first ever validated observation of WMN in human long-range WM tracts, laying the foundation for future research on the phenotype and function of these neurons, and how they may be affected in brain disorders.

## Introduction

In the mammalian brain, there are two structurally and functionally distinct compartments: the grey matter (GM) and the white matter (WM). The GM contains a high density of neuronal and glial cell bodies, cell processes and synapses (neuropil) as well as a low density of myelinated axons. The WM, on the other hand, is mainly composed of a high density of myelinated axons interspersed with glial cells. Sparse neurons also occur in the WM, more specifically in cortical WM. In 1867, Thomas Meynert first described the presence of neuronal cell bodies within the human cortical white matter (Meynert 1868). This was followed by decades of research attempting to classify and quantify these cells (for a complete history review see (Judaš et al. 2010)). First called “interstitial neurons” by Ramón y Cajal, they have since received many names, such as displaced, aberrant, heterotopic, paragriseal, intramedullary and intercalated neurons (Das and Kreutzberg 1969). Due to the inconsistent nomenclature of WM neurons we will be using the encompassing term “white matter neurons” (WMN) throughout this paper to describe neuronal cell bodies within human cerebral WM (following (García Marín et al. 2010a)).

Cortical WMN are generally divided into two separate categories: superficial and deep. Superficial WMN are neurons that are directly below layer VI of the GM and reside in the WM of the sulci and gyri of the cortex. Deep WMN reside further from the GM and are associated with long-range WM tracts (Fischer and Kukley 2024). There is currently no other, more precise way, to distinguish superficial from deep WMN. Some researchers have differentiated them based on the WM thickness of their sample (Eastwood and Harrison 2003), and others have arbitrarily set superficial WMN to be between 0 to 175μm from the GM/WM border and deep WMN to be between 375 to 550μm from the GM/WM border (García Marín et al. 2010a). The density of these neurons has been shown to decrease as the distance from the GM increases (Kostovic and Rakic 1990; Akbarian 1993; Beasley et al. 2002; Kubo 2020).

The origin of WMN is still under debate, as there are two differing opinions surrounding their development. Some researchers argue that the superficial WMN are derived from the subplate, a transient structure during development. The subplate is known to enable direct thalamocortical pathfinding and serve as a transient glutamatergic input for the developing cortex (Suarez-Sola et al. 2009). In humans, the subplate begins to slowly dissolve through apoptosis at the end of gestation and during the early postnatal period (Kostovic and Rakic 1990; Judaš et al. 2010). Some neurons persist postnatally and reside in the cortical and subcortical white matter, becoming WMN (Kostovic and Rakic 1990). Other researchers have argued that there is a constant supply of superficial WMN that are generated throughout later stages of corticogenesis, with the WMN being directly connected to WM with specific functions (Suarez-Sola et al. 2009). Furthermore, deep WMN are believed to be remnants of the subventricular zone neurons. These neurons are thought to be present in the fetal brain and differentiate into deep WMN (Kubo 2020; Fischer and Kukley 2024). Despite different developmental origins, very little is still known about which classes and subtypes of neurons survive and become WMN.

WMN have been phylogenetically conserved across rodents, cetaceans, and primates, highlighting a potential evolutionary significance (Mortazavi et al. 2016). The density of WMN is greater in humans and non-human primates (NHP), with lower densities observed in rodents. This could be due to the proportionally much smaller white matter volume and connections in rodent neocortex (Suarez-Sola et al. 2009). Alternatively, such lower densities may be due to different birthplace, migration and developmental pathways (Clancy 2009). When comparing NHP and humans, few differences in WMN morphology and density are seen in the frontal cortex (García Marín et al. 2010b; Raghanti et al. 2013; Swiegers et al. 2019). However, when analyzing the total number of WMN in NHP and humans, the WMN population consisted of 2.5% of total neurons in NHP (Swiegers et al. 2019), compared to 3.5% in humans (Sedmak and Judaš 2019). Altogether, the expansion of WMN highlights an evolutionary significance, with an increase density of WMN observed in higher cognitive order species. Though the exact function of WMN remains unknown, evidence suggests that these neurons are integrated into mature neural circuits and may play a role in regulating the flow of information in the cortex (Sedmak and Judaš 2021). Furthermore, they are posited to regulate cerebral blood flow due to their expression of nitric oxide and the close apposition of their axons to blood vessels (Okhotin and Kupriyanov 1997; Sedmak and Judaš 2021).

Due to the developmental and evolutionary importance of WMN, their potential implication has been investigated in several neuropsychiatric disorders. Particularly, WMN are believed to be important for the pathogenesis of schizophrenia. The density of cortical WMN has been found to be altered in patients with schizophrenia, providing evidence for the early developmental component of this psychopathology (Eastwood and Harrison 2005). For instance, in the superficial WM of the dorsolateral prefrontal cortex, the density of WMN was found to be increased in schizophrenia, highlighting the potential implication of these neurons in developmental abnormality (Eastwood and Harrison 2005). Alongside the dorsolateral prefrontal cortex, many other brain regions have shown significant changes in WMN densities in schizophrenia (Fischer and Kukley 2024). Similarly, other studies have determined an increased number of WMN in both superficial and deep WM in individuals with autism spectrum disorder (Fischer and Kukley 2024). Despite these findings, very little is known about WMN connectivity and functional significance throughout the brain.

In recent studies, WMN have been more systematically quantified to determine their densities across cortical WM. These have been determined in Brodmann areas 10, 17/18, 20 and 24, with the greatest density of WMN found in the frontal cortex (2, 660 ± 153.2 neurons/mm^3^) (García Marín et al. 2010b). Furthermore, Sedmak and Judaš classified WMN directly beneath the orbitofrontal cortex, cingulate cortex, dorsolateral prefrontal cortex, temporal cortex and occipital cortex (Sedmak and Judaš 2019). They also determined that the average number of WMN in the entire human brain is in the range of 450-670 million neurons, highlighting a significant and extensive population of neurons (Sedmak and Judaš 2019). In addition to these data, many neuropathological studies have examined the density of WMN in specific brain regions (Akbarian 1993; Anderson et al. 1996a; Kirkpatrick et al. 1999; Beasley et al. 2002; Rioux et al. 2003; Kirkpatrick et al. 2003; Eastwood and Harrison 2003; Connor et al. 2009; Yang et al. 2011; McFadden et al. 2016; Tsai et al. 2020). Neurons have also been observed in single nucleus RNA datasets of human WM (Jäkel et al. 2019). Despite the abundance of studies on WMN, none have sought to explore and characterize neurons in long-range WM tracts. Long-range WM tracts are essential for efficient communication between distant cortical areas, allowing for synchronous firing and overall brain coordination (Laughlin and Sejnowski 2003). To our knowledge, there have been no studies examining the presence of WMN in long-range WM tracts in the human brain.

Therefore, in this study, we investigated the presence of neuronal cell bodies within two WM tracts in humans: the uncinate fasciculus (UF) and corpus callosum (CC). The UF is a long-range association fiber that connects the orbitofrontal cortex with the anterior temporal lobe and has known implications in mnemonic-based associations and valence-based decision making (Von Der Heide et al. 2013). The CC is a commissural fiber that connects the left and right hemispheres of the brain and its function is to integrate and transfer sensory, motor and higher-level cognitive signals between the hemispheres (Roland et al. 2017).

In order to validate our methods, we also analyzed brain slices from the ventromedial prefrontal cortex (vmPFC). The prefrontal cortex has been highly studied with regards to cortical WMN, so using these density results from the literature (Anderson et al. 1996b; Eastwood and Harrison 2005; García Marín et al. 2010b) allowed us to set it as a reference point. Since both the UF and CC are long-range WM tracts and contain only deep WMN, we only investigated and quantified deep WMN (according to the values set by (García Marín et al. 2010b)) in the vmPFC in order to allow for direct comparisons.

We used histological techniques including immunohistochemistry (IHC) and fluorescence in situ hybridization (FISH) using RNAScope®, in order to determine the presence and density of WMN within the UF and CC. We also used neuronal subtype markers to determine the proportion of excitatory and inhibitory neurons. Finally, we aimed to determine whether there were significant differences in neuronal densities between these WM tracts, and compared to cortical WM.

## Materials and Methods

### Human post-mortem brain tissue

Brain samples were provided by the Douglas-Bell Canada Brain Bank (DBCBB; Montréal, Canada). Brains were obtained by the DBCBB thanks to a special agreement with Québec Coroner’s Office, with informed consent from next of kin. Phenotypic information was collected with standardized psychological autopsies, as described previously (Dumais et al. 2005; Perlman et al. 2021). We investigated only “control” subjects, with no known neurological or psychiatric disorders, as supported by coroner records and clinical files when available. Toxicological assessments and medication prescription data was also collected. We selected subjects with a wide age-range and included both males and females.

All dissections were performed on fresh-frozen WM from the left hemisphere, by the expert brain bank staff, with the guidance of a human brain atlas (Mai et al. 2008). The UF was dissected at Brodmann area 38 (temporal pole), which is an area that primarily contains UF fibers, without concurrent fibers from other tracts (Von Der Heide et al. 2013). This region constitutes the temporal terminations of the UF fibers. A second UF region was dissected at the junction of the frontal lobe and the temporal lobe laterally to the amygdala, approximately - 8.0mm from the center of the anterior commissure (n=4). The CC was dissected at the level of the genu, which forms the anterior portion of the CC, along with the rostrum (Kier and Truwit 1996). Both CC and UF samples were from the same 12 subjects (Table 1). The vmPFC was dissected at the level of the rectus gyrus, medial or intermediate orbital gyri, approximately 48mm from the center of the anterior commissure (n=7) (Table 1). Samples were stored at −80°C until use.

**Table 1.** Fresh frozen human brain sample characteristics. Data presented as mean ± standard deviation. M/F: male/female, PMI: postmortem interval

|  | CC/UF | vmPFC |
| --- | --- | --- |
| N | 12 | 7 |
| M/F | 9/3 | 4/3 |
| Age | 48.08 ± 18.04 | 58.86 ± 11.20 |
| PMI (h) | 49.18 ± 29.06 | 57.43 ± 17.61 |
| pH | 6.38 ± 0.35 | 6.12 ± 0.24 |

### Histology

The frozen UF, CC and vmPFC samples were cut serially using a Leica CM1950 cryostat to a thickness of 10μm, collected on Fisherbrand™ Superfrost™ Plus Microscope Slides (reference 12-550-15) and stored at −80 °C. A cresyl violet stain was performed to confirm the presence of WM within the cryosectioned tissue. Slides were left to warm up to room temperature and then incubated in 0.1% Cresyl Violet for 15 min. Then, they were dehydrated using successive ethanol baths of increasing concentrations, followed by 5 min in xylenes. The slides were coverslipped using Permount mounting medium. This allowed us to identify any possible GM contamination within the brain tissue. Once we determined the clear areas of WM, we took the immediately adjacent brain sections for each subject to perform the in situ hybridization and IHC experiments.

UF and CC samples surrounded with large amounts of contaminating GM were excluded to avoid any possible confusion with cortical WM. This exclusion criterion led to one UF sample to be excluded from both FISH and IHC analyses. In addition, there was one fewer vmPFC subject in the IHC results compared to the FISH results due to limited tissue availability.

### Fluorescence in situ hybridization (FISH)

All in situ hybridizations were performed using Advanced Cell Diagnostics RNAscope® probes and reagents following the manufacturer’s instructions. The slides were fixed in 10% formalin for 15 min at 4°C, followed by successive ethanol dehydration steps with increasing concentrations of ethanol. To remove endogenous peroxidase activity, the brain tissue was incubated in 3% hydrogen peroxide for 10 min at room temperature followed by probe hybridization for 2 hours at 40°C. The following probes were used: Hs-SLC17A7-C1 (catalog number 415611), Hs-RBFOX3-C2 (catalog number 415591-C2) and Hs-GAD1-O1-C3 (catalog number 573061-C3). Amplifiers were used to increase the signal, followed by probe-specific HRP detection. Opal dyes with tyramide signal amplification (TSA) were used to achieve visual signal through HRP binding, resulting in colour-specific channels for each of the probes. Opal 520 (Akoya Biosciences FP1487001KT), Opal 570 (Akoya Biosciences FP1488001KT) and Opal 690 (Akoya Biosciences FP1497001KT) were all diluted 1:300. To quench endogenous autofluorescence, 5% TrueBlack in 70% ethanol was used for 30 seconds. To coverslip, Vectashield Vibrance mounting medium with 4′,6-diamidino-2-phenylindole (DAPI) (Vector Laboratories H-1800-10) was used, which labelled the cell nuclei. The slides were kept at 4°C until imaging.

### Immunohistochemistry (IHC)

The slides were fixed for 10 min at room temperature, followed by three washes using phosphate-buffered saline (PBS). Endogenous peroxidase activity was quenched using 3% hydrogen peroxide for 10 min, followed by a wash in PBS. To prevent non-specific binding, the samples were blocked with 2% natural horse serum in PBS+0.2% Triton X-100 for 1 hour at room temperature. NeuN mouse primary antibody (Millipore MAB377) was added at 1:500 to the blocking solution and incubated at 4°C overnight. After washing, the samples were incubated for 30 min at room temperature using biotinylated horse anti-mouse secondary antibody (Vector Laboratories BA-2000-1.5) at 1:1000. Using the Vectastain Elite ABC kit (Vector Laboratories PK-6100), the streptavidin conjugated horse radish peroxidase (HRP) recognizes the biotinylated secondary antibody. The HRP then reacts with the chromogenic 3,3’-diaminobenzidine substrate (DAB; Vector Laboratories SK-4100), resulting in in a dark brown precipitate in the cells containing the NeuN antigen. Sections were incubated in the DAB solution for 20 min, followed by a wash in distilled water to stop the reaction. Finally, the samples were dehydrated using successive ethanol washes of increasing concentrations, followed by coverslipping with permount mounting media.

### Image Analysis

Sample images were taken using an Evident Scientific FV1200 confocal microscope at 40x magnification. Full slide images were taken using an Evident Scientific VS120 Slide Scanner at 20x and analyzed using QuPath (v.0.5.1). To determine the WMN density and proportion, 6 region of interest (ROI) boxes with an area of 1.875 mm^2^ were randomly placed across the WM of each sample. For both FISH and IHC, NeuN was used in order to determine the GM/WM border. Due to the marked decrease in neuronal density after layer VI, it was possible to ensure that all neurons counted were in the WM. Furthermore, all boxes were placed at minimum 375μm away from the GM/WM border in order to ensure that all neurons counted were not superficial WMN (following (García Marín et al. 2010b)). Since all neurons in long-range WM tracts would be deep WMN, this ensured that we were comparing solely deep WMN densities between the vmPFC, UF and CC.

For RNAScope, image brightness settings were adjusted on QuPath, using both a positive SLC17A7 neuron and GAD1 neuron as a reference point in order to ensure that all background fluorescence was removed. Due to the paucity of reference neurons in the CC, the settings for the UF for each subject were applied to the CC. For a neuron to be considered SLC17A7 positive (+), it had to contain at least 3 positive puncta and colocalize with DAPI. For GAD1, signal stronger than the background that colocalized with DAPI constituted a GAD1 positive (+) neuron.

For IHC, the images were also analyzed on QuPath. Brightness settings were adjusted to achieve the highest contrast between positive DAB-labelled WMN and background. All cells were counted as NeuN positive (+) if they presented the attributes of a cell body and contained a strong DAB labeling. For both the IHC and FISH, all boxes were manually counted blindly by one experimenter to ensure consistency.

### Statistics

Statistical analysis was conducted using R (version 4.4.2). Linear modelling with the lmer package was applied for all analyses, with brain region, age, sex, postmortem interval (PMI), and pH as factors, with subject ID as a repeated measure. Tukey’s HSD was implemented with the emmeans package for post-hoc testing. Pearson correlations were used to compare values across RNAScope® and IHC. P-values < 0.05 were considered significant. All data are presented as mean ± standard error of the mean unless otherwise specified.

## Results

We observed WMN in all three regions, with varying densities. All vmPFC subjects contained WMN in all ROI boxes counted. In the CC of several subjects, WMN were absent in all of the sampling boxes counted. To ensure that the WMN counted in the CC were not artefacts, we separately recounted the CC subjects to confirm that our results were accurate. There was only one subject that had no WMN in the UF.

### Presence of WMN validated at the RNA level

Using RNAScope® for FISH, we were able to label excitatory and inhibitory neurons expressing SLC17A7 and GAD1, respectively. A representative image example image of the UF is illustrated in Figure 1A, while the CC and vmPFC can be found in Supplementary Fig. 1. We observed meaningful quantities of WMN in both the vmPFC and UF. There were much fewer quantities of WMN in the CC. Notably, there was a significantly higher density of SLC17A7+ neurons in the vmPFC compared to the CC (mean vmPFC: 10.2 ± 1.35 neurons/mm^2^; mean CC: 0.18 ± 0.079 neurons/mm^2^; *p*<0.0001, Fig. 1B), and in the UF vs the CC (mean UF: 6.92 ± 0.63 neurons/mm^2^; mean CC: 0.18 ± 0.079 neurons/mm^2^; *p*=0.0003, Fig. 1B). There were more SLC17A7+ neurons in the vmPFC compared to the UF, however this difference was not significant (*p*=0.231, Fig. 1B). For GAD1+ neurons, there was a significantly higher density in vmPFC vs CC (mean vmPFC: 2.59 ± 0.38 neurons/mm^2^; mean CC: 0.24 ± 0.057 neurons/mm^2^; *p*=0.0260, Fig. 1B) UF vs CC (mean UF: 2.54 ± 0.29 neurons/mm^2^; mean CC: 0.24 ± 0.057 neurons/mm^2^; *p*=0.0018, Fig. 1B), but there was no difference between the UF and vmPFC (*p*=0.995, Figure 1B). The covariates (age, sex, PMI and pH) did not show significance in either model. We next determined the ratio of excitatory to inhibitory neurons by dividing the density of SLC17A7+ cells by the density of GAD1+ cells. The CC had the lowest ratio of excitatory to inhibitory neurons (0.73 ± 0.36), followed by the UF (2.73 ± 0.40), with the vmPFC having the highest ratio (3.93 ± 0.78) (Figure 1C). NeuN labeling with FISH gave rise to a high level of background signal, making us unable to quantify WMN that were both NeuN+ as well as SLC17A7- and GAD1-negative. However, NeuN did provide a robust way to determine the WM/GM border and validate SLC17A7+ and GAD1+ cells, but we could not use it for further classification due to the important background.

**Figure 1.**
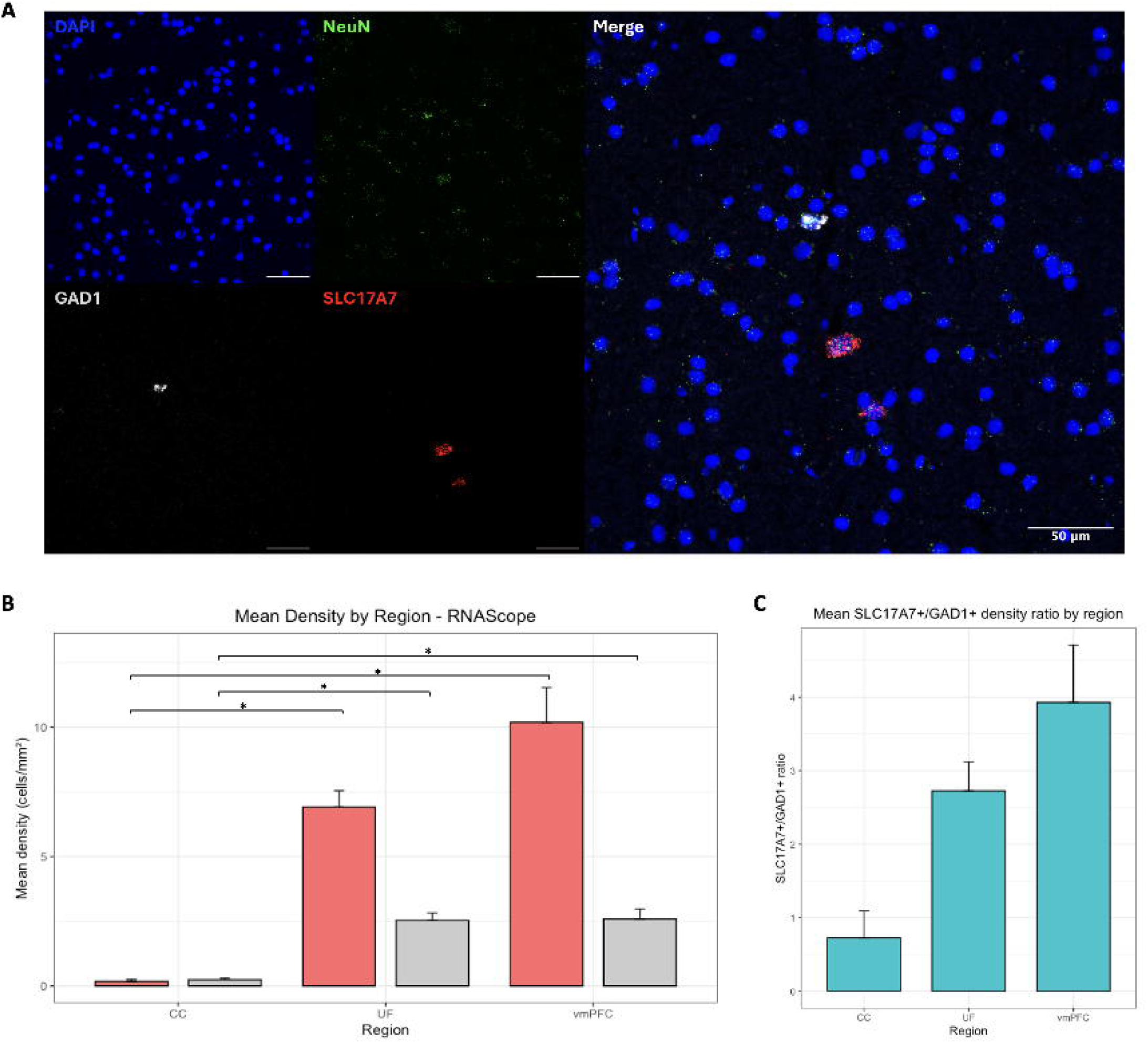
White matter neurons detected via RNA level expression show significant regional differences in neuronal densities. **A)** Images of white matter neurons in the UF using RNAScope for NeuN (green), GAD1 (white) and SLC17A7 (red), with nuclei stained using DAPI. Imaged on an Evident Scientific FV1200 confocal at 40X magnification. Scale bar=50μm. **B)** White matter neuron densities for both excitatory (red) and inhibitory (grey) across regions. **C)** Proportion of excitatory to inhibitory white matter neuron densities. vmPFC: ventromedial prefrontal cortex, UF: uncinate fasciculus, CC: corpus callosum

### Presence of WMN validated at the protein level

IHC was used to label the pan-neuronal marker NeuN at the protein level, and WMN were observed in all three brain regions (Fig. 2). When comparing the three brain regions, there was a significantly higher WMN density in the vmPFC compared to the CC (mean vmPFC: 12.3 ± 1.15 neurons/mm^2^; mean CC: 0.29 ± 0.11 neurons/mm^2^; *p*<0.001, Figure 2A) and in the UF compared to the CC (mean UF: 10.7 ± 0.85 neurons/mm^2^; mean CC: 0.29 ± 0.11 neurons/mm^2^; *p*<0.001, Figure 2A). There was no difference between the UF and vmPFC (*p*=0.971, Figure 2A). Similar to the FISH results, none of the model covariates (age, sex, PMI, and pH) showed any significance.

**Figure 2.**
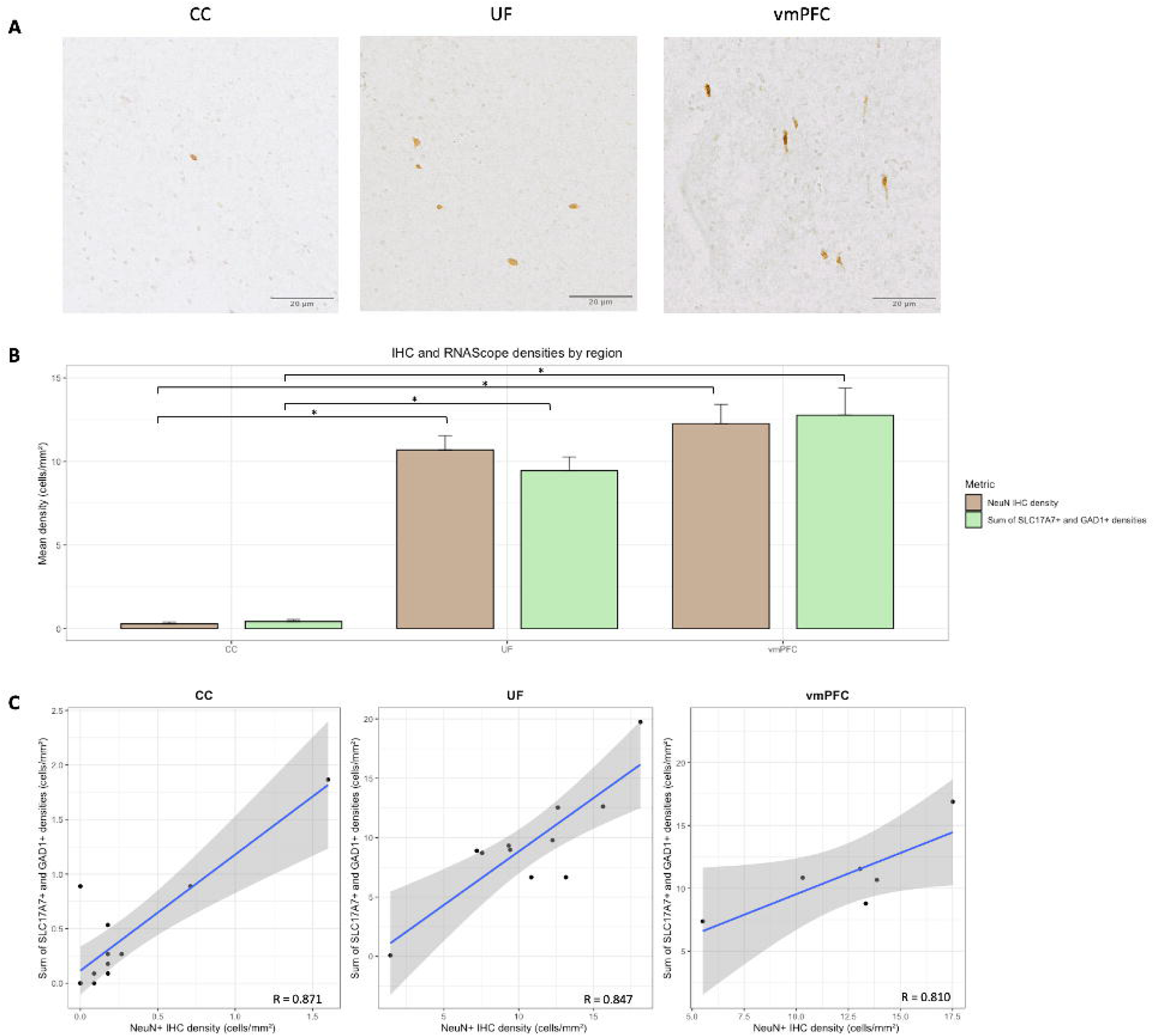
White matter neurons are detected at the protein level. **A)** Images of DAB-stained white matter neurons in all brain regions using immunohistochemistry. Imaged on an Evident Scientific VS120 Slide Scanner at 40x. Scale bar=20μm. **B)** Neuronal densities for both immunohistochemistry (brown) and the sum of both excitatory and inhibitory neurons for the RNAScope (green). For both the immunohistochemistry and RNAScope, white matter neuron densities significantly increased between the CC and vmPFC and the CC and UF. No significant changes were observed between the UF and vmPFC. **C)** Strong correlation between immunohistochemistry and RNAScope white matter neuron densities. CC: corpus callosum, UF: uncinate fasciculus, vmPFC: ventromedial prefrontal cortex.

### Comparing RNA and protein-level expression

By adding up the total number of neurons counted following FISH labeling, it was possible to compare WMN densities counted following FISH and IHC. When summing the excitatory and inhibitory neurons together, the vmPFC density was significantly higher than the CC density (mean vmPFC: 12.8 ± 1.61 neurons/mm^2^; mean CC: 0.42 ± 0.12 neurons/mm^2^; *p*=0.0001, Figure 2B) and the UF density was significantly higher than the CC density (mean UF: 9.45 ± 0.82 neurons/mm^2^; mean CC: 0.42 ± 0.12 neurons/mm^2^; *p*=0.0003, Figure 2B). The vmPFC density was higher than the UF density, though not significantly (p=0.513). Again, the covariates (age, sex, PMI and pH) did not show significance in the model. As expected, the densities following FISH and IHC labelings were similar within each region (Fig. 2B). As such, we found that the densities from both the RNA- and protein-stained tissues were strongly correlated for the CC (R=0.871, Fig. 2C), the UF (R=0.847, Fig. 2C), and the vmPFC (R=0.810, Fig. 2C).

### Further validation of UF WMN densities

Due to our observation of substantially more WMN in the UF than in the CC (22.5 times more, according to the sum of the FISH markers), we sought to validate whether these WMN were limited to the terminations of the temporal segment of the UF (dissected at BA38) or if they were present in similar quantities elsewhere in the tract. To this end, we performed the same FISH and IF protocols on sections (n=4) of the UF dissected from a second location, specifically at the junction of the frontal lobe and the temporal lobe lateral to the amygdala. This specific region was chosen because it was one of the few areas from which the UF can be clearly distinguished from other proximal tracts. Fig. 3A illustrates the two different UF regions (with the original UF dissection denoted as UF_a_ and this new dissection denoted as UF_b_). The 4 sections analyzed in UF_b_ were derived from a subset of subjects from the UF_a_ cohort (Supplementary Table 1). UF_b_ SLC17A7+ and GAD1+ FISH densities did not show significant differences from UF_a_ (SLC17A7 *p* = 0.348, GAD1 *p* = 0.396, Fig. 3B). The ratios of excitatory to inhibitory WMN in both UF locations are depicted in Fig. 3C. Additionally, the sum of SLC17A7+ and GAD1+ does not show significant differences between the two UF locations (*p* = 0.381, Fig. 3D). At the protein level, NeuN+ cells were also present in UF_b_, with no significant difference in density from UF_a_ (NeuN+ *p* = 0.560, Fig. 3D). The RNA- and protein-level measurements correlate strongly in UF_b_ (R = 0.903, Figure 3E). None of the covariates (age, sex, PMI, and pH) showed significance in any of the models. In summary, these experiments show that the substantial presence of WMN is not limited to the UF terminal fibers but is present throughout the UF temporal segment.

**Figure 3.**
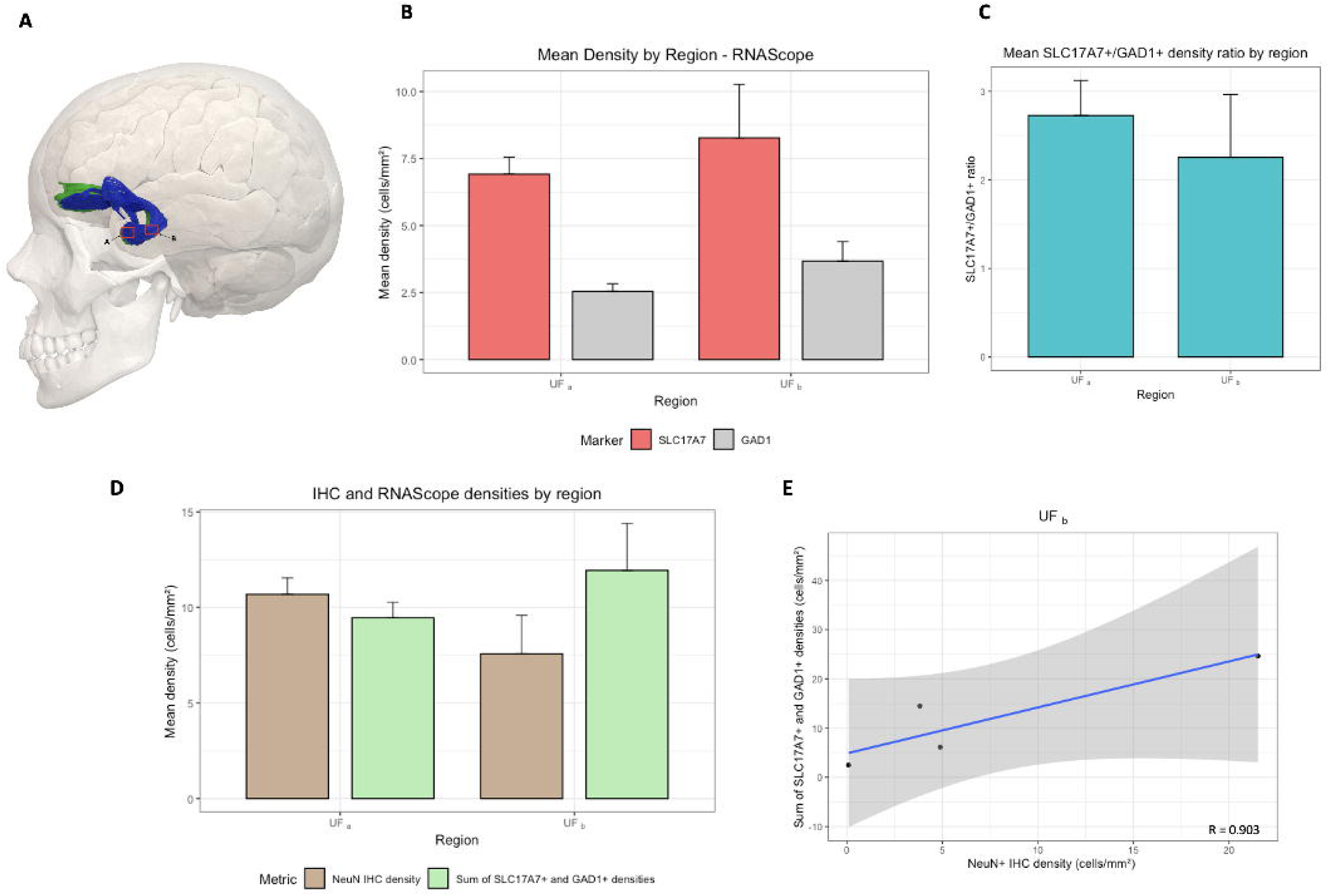
White matter neurons validated at a second location along the UF. **A)** Schematic demonstrating the UF dissection locations. The left UF is depicted in blue and the right UF is depicted in green. Both dissections were performed in the left hemisphere. Box A corresponds to UF_a_ dissected at BA38, and box B corresponds to UF_b_ dissected at the junction of the frontal lobe and the temporal lobe lateral to the amygdala. The image was adapted from <u>“BodyParts3D,© The Database Center for Life Science licensed under CC Attribution-Share Alike 2.1 Japan.”</u> **B)** White matter neuron densities for both excitatory (red) and inhibitory (grey) across both UF locations. **C)** Proportion of excitatory to inhibitory white matter neuron densities across both UF locations. **D)** Neuronal densities for both immunohistochemistry (brown) and the sum of both excitatory and inhibitory neurons for the RNAScope (green) across both UF locations. **E)** Strong correlation between immunohistochemistry and RNAScope white matter neuron densities in UF_b_ (R = 0.903). CC: corpus callosum, UF: uncinate fasciculus, vmPFC: ventromedial prefrontal cortex

## Discussion

In this study, we investigated the density and phenotype of WMN in two human long-range WM tracts. We found that WMN were present in both tracts, albeit in lower densities than in cortical WM (vmPFC). The density of WMN in the UF was not statistically different from WMN density in the vmPFC, but was descriptively consistently lower, particularly for SLC17A7 cells. The CC had the lowest density of WMN, and contained very few neurons. Overall, these results indicate that there are neuronal cell bodies within at least two human long-range WM tracts, with varying densities.

Due to the new approaches used in this study, we aimed to validate our methods by comparing our measurements of vmPFC WMN (internal control region) with existing measurements in the literature. We found that the density of deep WMN in the vmPFC was approximately 12.5 neurons/mm^2^. This corresponds to results previously described in other cortical WM areas. In the dorsolateral prefrontal cortex, densities of deep WMN have been reported to vary between 9-12 neurons/mm^2^ (Eastwood and Harrison 2005; Yang et al. 2011). Similarly, in the parahippocampal gyrus, a density of 9 neurons/mm^2^ has been reported for deep WMN (Eastwood and Harrison 2005). Our results for vmPFC WMN densities are therefore in the same range as those reported in previous studies, adding further confidence to the reliability of our observations and density measurements in the UF and CC.

Our results illustrate that WMN contain populations of both excitatory (glutamatergic) and inhibitory (GABAergic) neuronal subtypes. To date, the presence of glutamatergic WMN has been unclear, as one of the common glutamatergic marker used has been SMI32, which can also label GABAergic neurons (Kelly et al. 2019; Fischer and Kukley 2024). In a review article by Fischer and Kukley (2024), it was suggested that WMN could be capable of co-releasing GABA and glutamate, a mechanisms identified elsewhere in the brain (Wallace and Sabatini 2023; Ceballos et al. 2024). Our results indicate that this is not the case as we did not find any WMN that co-expressed both SLC17A7 and GAD1, indicating that WMN do not co-release glutamate and GABA in the regions we considered. When considering the proportion of excitatory to inhibitory WMN, there was a lower ratio in the UF compared to the vmPFC. In the UF, inhibitory neurons accounted for approximately 25% of all WMN counted, whereas in the vmPFC, inhibitory neurons accounted for around 20%. This suggests a potential regional difference for WMN subtypes. These results are aligned with the literature, both brain-wide and specifically for WMN (García Marín et al. 2010b; Alreja et al. 2022). Across the whole human brain, approximately 10-25% of all neurons are inhibitory (Alreja et al. 2022). When looking specifically at WMN, it has been reported that around 25% of all WMN in the human neocortex express inhibitory markers (García Marín et al. 2010b).

In recent literature, increasing evidence has suggested that levels of RNA and protein are not as correlated as originally thought (De Sousa Abreu et al. 2009; Koussounadis et al. 2015). This highlights the importance of studying RNA and protein expression in a side-by-side comparison, rather than individually. Our research confirmed and correlated the density of WMN at both the RNA and protein-level, providing robust evidence for the presence of these neurons in WM tracts.

By comparing the two long-range WM tracts analyzed in this study, it is evident that the CC contains a much smaller quantity of WMN compared to the UF. Although there have been no human postmortem studies on CC WMN, there have been studies on mice and rats confirming the presence of WMN in the CC, but neither of these studies determined the WMN density (Von Engelhardt et al. 2011; Barbaresi et al. 2014). These rodent results are aligned with our human postmortem investigation and support the notion that WMN in WM tracts might be a feature present in different mammalian species. One potential reasoning for the low WMN density in the CC in our study is that the CC was dissected at the genu, which is anatomically further from the GM, and it is known that the density of WMN progressively decreases as the distance from the GM increases (Kubo 2020). As for the UF, UFa was dissected at Brodmann area 38, which is adjacent to the temporal pole and thus anatomically close to GM. The UF_b_ dissection likely intersected a part of the claustrum. As such, both UF dissections were spatially more proximal to GM than the CC. This could potentially explain the sharp decrease in WMN in the CC and highlights that specific regional WMN densities may be related with the distance to GM.

A limitation of this study was the inability to classify the WMN density in terms of distance from the GM. Most of our tissue samples contained purely WM, meaning we could not determine the distance from the GM, as recommended by previous literature (Sedmak and Judaš 2019). Furthermore, it has been reported that specific subsets of neurons are NeuN-negative, such as cerebellar Purkinje cells, Cajal-Retzius cells of the neocortex, mitral cells in the olfactory bulb and cerebellar interneurons (Kumar and Buckmaster 2007). As such, using NeuN as a marker for all neurons may constitute a limitation. However, these neuron subtypes are not found in the brain regions examined in this study and are therefore unlikely to impact the WMN densities observed here. It has also been shown that NeuN labels the overall general population of mature WMN, whereas all other common neuronal markers only label a small subset, making it the most robust marker for WMN (Duchatel et al. 2019).

There have been reports of peptidergic WMN being found in human cortical and subcortical WM (Ang and Shul 1995; Wiesner et al. 2024). Depending on the immunochemical marker used, Ang and Shul found that the density of peptidergic WMN varied from 0.89-4.55 neurons/mm^2^ (Ang and Shul 1995). These authors did not report the distance from the GM so it is unclear whether these peptidergic WMN reside in both the superficial and deep WM. One study has shown that a proportion of GABAergic WMN also express the serotonin receptor 5-HT_3A_ (Von Engelhardt et al. 2011). Therefore, an important future direction would be to further characterize WMN subtypes by exploring potential peptidergic phenotypes.

To date, researchers have investigated cortical WMN primarily in Alzheimer’s disease and schizophrenia (for a complete review see (Fischer and Kukley 2024)). As such, performing case-control studies evaluating WMN across different brain regions in various neuropsychiatric disorders is another important future direction. Finally, a crucial next step is to investigate the relationship between GM proximity and WMN density in long-range WM tracts.

To conclude, this study represents the first ever characterization of WMN in human long-range WM tracts. Our study provides robust evidence for the presence of WMN, of both excitatory and inhibitory subtypes, in long-range WM tracts (CC and UF). The CC contained an extremely low density of WMN, with the UF containing a significantly higher density, in the range similar to that measured in the vmPFC WM. These results were confirmed at both the RNA and protein-level and correlated with each other, confirming the reliability of our quantification methods. The UF WMN results were further validated in a second region along the UF temporal segment. Regional differences were observed, both in WMN density and subtype proportions, highlighting the importance of future research on different brain regions. Our results lay a strong foundation for future research into the molecular, functional, and pathological role of this understudied neuronal population.

## Acknowledgements

We would like to thank the Douglas-Bell Canada Brain Bank staff and Bita Khadivjam at the Douglas Molecular and Cellular Microscopy Platform. Most importantly, we would like to thank the brain donors and their next of kin who consented to donating the brains of their loved ones.

