## Supplementary Material for "Characterizing neuronal cell bodies in human postmortem cerebral white matter tracts"

|  | UF_a_ | UF_b_ |
| --- | --- | --- |
| N | 12 | 4 |
| M/F | 9/3 | 4/0 |
| Age | 48.08 ± 18.04 | 36.50 ± 16.30 |
| PMI (h) | 49.18 ± 29.06 | 35.63 ± 41.48 |
| pH | 6.38 ± 0.35 | 6.53 ± 0.23 |

**Supplementary Table 1**

**Fresh frozen human brain sample characteristics for UF dissections**. Data presented as mean ± standard deviation. M/F: male/female, PMI: postmortem interval

**Supplementary Figure 1**

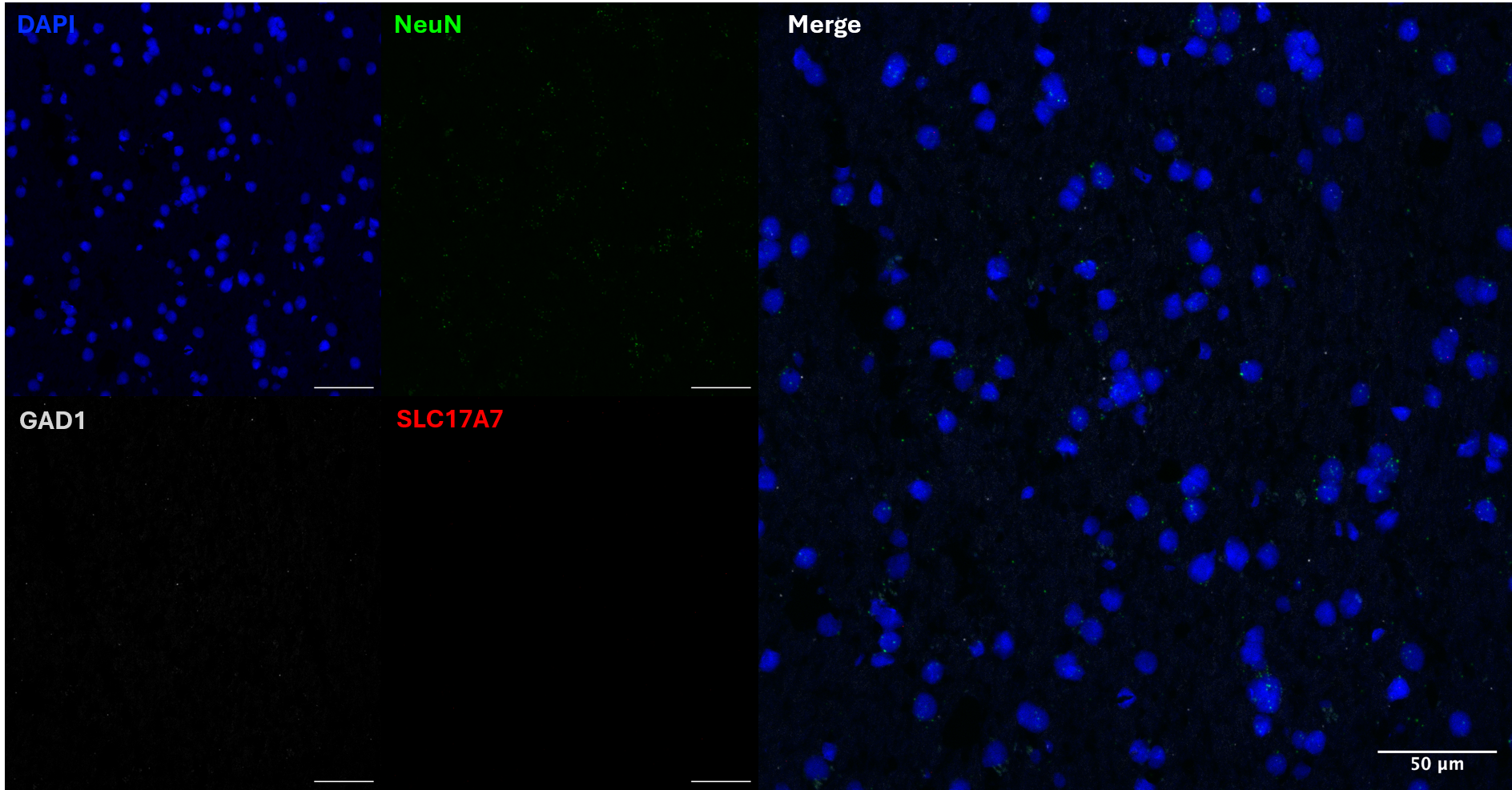

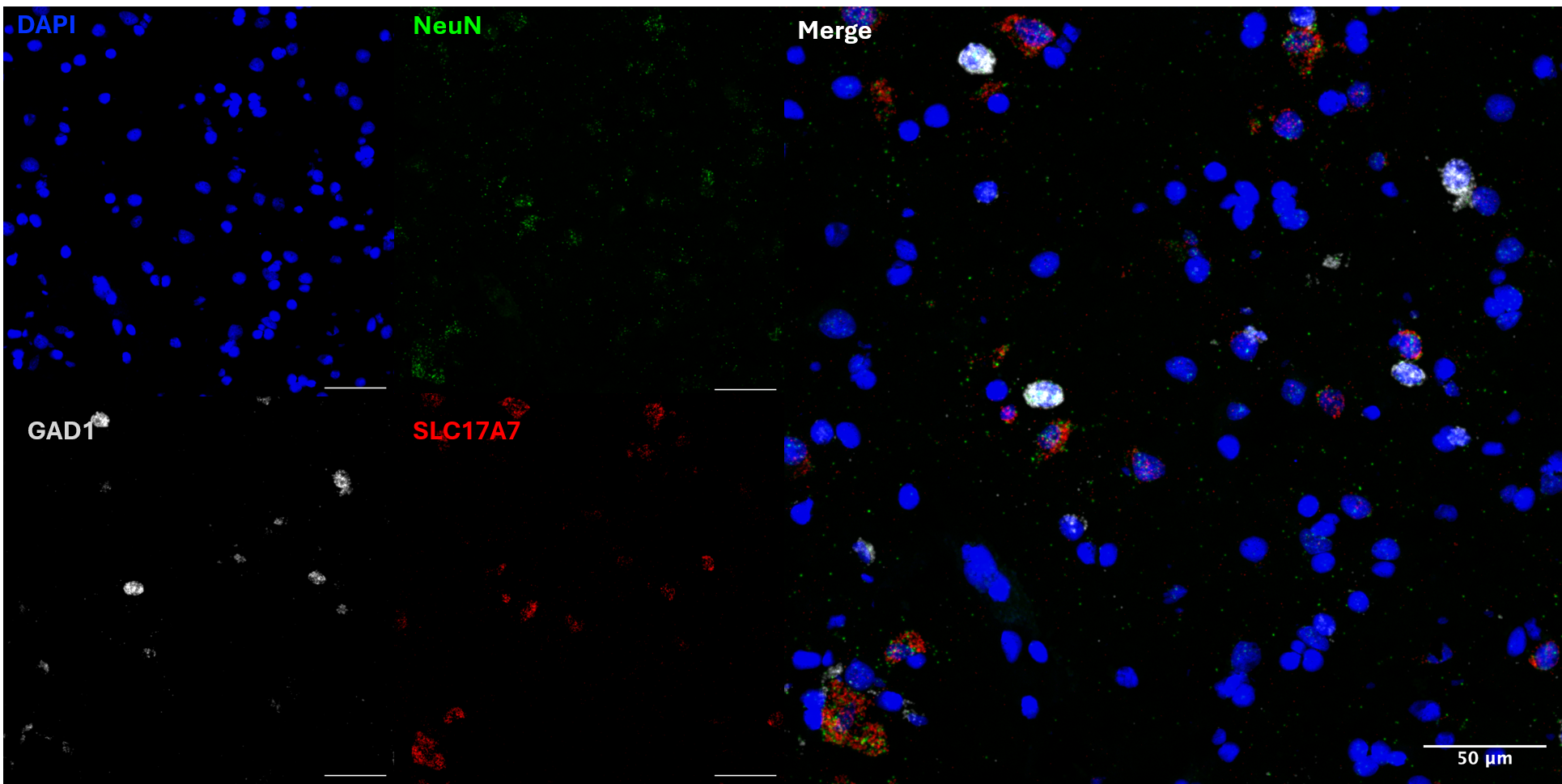

**A**

vmPFC

CC

**B**

Supplementary Figure 1. **Example images of white matter neurons using RNAScope.** The probes used are NeuN (green), GAD1 (white) and SLC17A7 (red), with nuclei stained using DAPI in the **A)** CC and **B)** vmPFC. There are no visible neurons in this section of the CC. Imaged on an Evident Scientific FV1200 confocal at 40X magnification. Scale bar=50μm. CC: corpus callosum, vmPFC: ventromedial prefrontal cortex.
